# Eccentricity, but not category, organizes infant ventral temporal cortex

**DOI:** 10.64898/2026.10.01.755396

**Authors:** Sarah Shi Tung, Emily M. Chen, Cleo Tay, Kylie Szymanski, Cameron Thomas Ellis

**Affiliations:** Department of Psychology, Stanford University, United States; Department of Psychology, Yale University, United States

**Keywords:** awake infant fMRI, ventral temporal cortex, retinotopic organization, category selectivity, fovea, periphery

## Abstract

Scene- and word-selective regions in adult ventral temporal cortex occupy consistent locations across individuals and align with eccentricity biases for peripheral versus central parts of the visual field, respectively. This relationship may reflect a developmental sequence in which initial preferences for visual field location influence the emergence of category-selectivity in these regions, driven by where each category appears in the visual field. To test this account, we used fMRI in awake, behaving infants aged 2–10 months, independently manipulating stimulus size (40° versus 6°) and category (scenes versus letters). We found eccentricity gradients across early visual cortex and ventral temporal cortex. Crucially, the collateral sulcus — a scene-selective, peripherally-biased region in adults — showed greater responses to big over small stimuli in infants, but no reliable preference for scenes over letters. These findings suggest that an eccentricity bias could provide a scaffold for later category specialization.

## INTRODUCTION

How the mature organization of the visual system emerges is a key question in developmental cognitive neuroscience. Two important properties of the adult primate visual system are retinotopic organization and category selectivity. Retinotopic organization refers to the systematic representation of visual field location across cortex^1^. One component of retinotopic organization is eccentricity, which organizes visual-field representations from the center of gaze to the periphery. Category selectivity refers to stereotypically-localized patches of cortex that respond selectively to ecologically relevant categories such as faces, scenes, and words^2–4^. In ventral temporal cortex, category selectivity is systematically coupled with eccentricity: regions preferring categories that are typically viewed centrally, such as faces and words, show foveal eccentricity biases, whereas scene-selective regions show peripheral eccentricity biases^5–8^. Understanding how eccentricity organization and category selectivity emerge in relation to one another requires studying the developing brain.

Both eccentricity organization and category selectivity seem to emerge remarkably early in visual development. Human infants possess spatial frequency maps (a corollary to eccentricity maps^9–14^) in early visual cortex by 5 months^15^ and category selectivity as early as 2 months^16–18^. The presence of both properties in infancy might suggest that eccentricity and category organization arise through separate mechanisms^19^. However, the proto-organization hypothesis^7,9,20,21^ offers an alternative account in which category-selective organization develops from earlier biases for lower-level visual features that co-occur with particular categories. Consistent with this hypothesis, work in neonatal macaques shows that eccentricity-related organization within ventral temporal cortex is present well before adult-like category selectivity emerges^9^. Hence, regions initially biased toward foveal or peripheral input may become tuned during development for categories typically processed in those portions of the visual field, such as letters in the fovea or scenes in the periphery. On this account, an eccentricity bias in ventral temporal cortex may shape the visual information that regions initially process, providing a scaffold for subsequent experience-dependent cortical specialization.

Here, we investigated the development of eccentricity and category responses using fMRI with awake, behaving infants (2-10 months). Given that infants cannot maintain fixation reliably, we designed a free-viewing paradigm to test eccentricity and category effects. We independently manipulated size (big versus small) and category (scenes versus letters) in a block design (Figure 1). We focused on the collateral sulcus (CoS), which is scene-selective and peripherally biased in adults^5–7^. We preregistered our hypothesis that an eccentricity bias would be present throughout infancy, whereas a category preference for scenes over letters would be weaker and emerge only in older infants, consistent with the proto-organization hypothesis.

**Figure 1.**
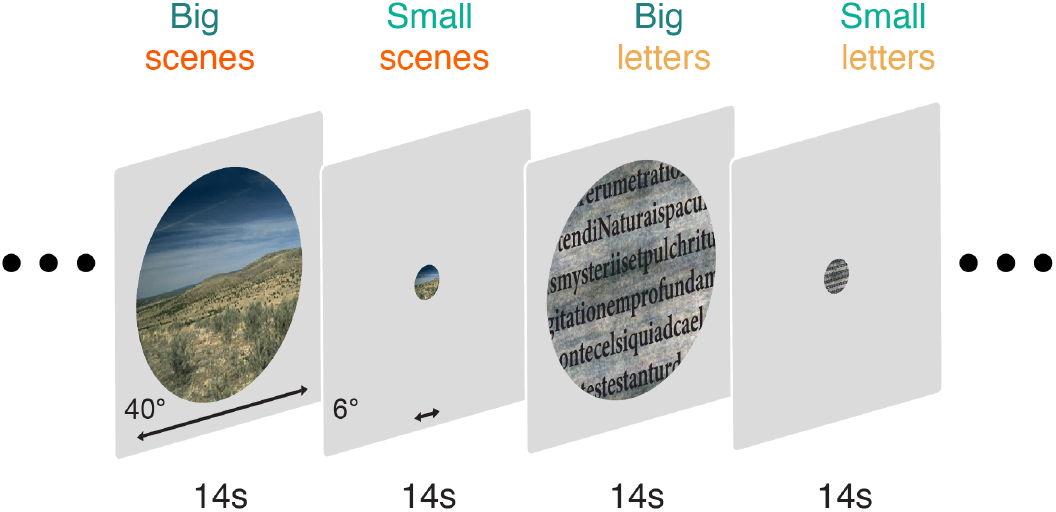
Experimental design. Participants viewed scene and letter stimuli at two sizes: big (40°) and small (6°). The same sequence of scene images shown in the scene conditions were repeated for the letter conditions, except that the scenes were phase-scrambled and in the background. Each condition lasted 14 s with 14 stimuli shown per condition. At the end of a set of all four conditions, a 6 s blank period was shown.

## RESULTS

We collected a preregistered sample of 24 infant sessions (2.1-9.6 months old) in which infants viewed blocks of scenes and letters, presented both big and small, yielding four stimulus conditions: big scenes, small scenes, big letters, and small letters (Figure 1). For comparison, we also collected an adult sample (N = 7) to validate our paradigm, which we report in the supplementary materials. Each condition was shown for 14 s. We performed our main analyses as specified by our preregistration and label additional analyses as exploratory.

### Looking Behavior

We manually coded infants’ gaze from video recordings collected during the experiment and asked whether looking behavior differed by stimulus size or category. Because infants freely viewed the stimuli rather than being required to maintain fixation, we compared the proportion of looking time directed to the center of the screen across conditions. In an exploratory analysis, we found a difference in looking behavior for stimulus size, in which infants looked more toward the center of the display during presentation of the small stimuli than the big stimuli (Figure S1A), as expected. Importantly, we found no difference between big and small stimuli for on-screen looking (Figure S1C), and no difference between scene and letter stimuli for center-looking (Figure S1B) or on-screen looking (Figure S1D). These results suggest that the stimulus conditions were similarly engaging overall, while the difference in center-looking between small and big stimuli reflected their spatial extent on the screen.

### Evidence of eccentricity gradients in infant visual cortex

We asked whether an eccentricity bias is present in infant early visual cortex (EVC). We used a GLM to estimate blood-oxygenation-level-dependent (BOLD) responses to the four stimulus conditions, and computed the contrast of all big versus all small stimuli (i.e., the size contrast). Because our big stimuli extended farther into the periphery, a reliable difference in the response to big versus small stimuli demonstrates evidence of an eccentricity bias.

We used the Wang probabilistic atlas^22^ to track eccentricity biases within early visual areas V1, V2, and V3 on the dorsal and ventral surfaces. Within each area, we sampled responses along the posterior-to-anterior axis (corresponding to the foveal-to-peripheral axis) (Figure 2A), and compared the mean size contrast at the posterior quartile with the mean size contrast at the anterior quartile, averaging across dorsal and ventral cortex. We found a significant increase in the size contrast, such that big stimuli evoked a stronger response than small stimuli in the anterior portion of all visual areas tested, relative to the posterior portion (Figure 2C): V1 (CI = [1.29, 2.66], d = 1.11, p *<* 0.001), V2 (CI = [1.15, 2.40], d = 1.10, p *<* 0.001), and V3 (CI = [0.97, 1.95], d = 1.13, p *<* 0.001). This is consistent with our preregistered prediction that infant EVC would show an eccentricity gradient. The same pattern of results was observed in our adult sample (Figure S2). Thus, infant EVC already shows an eccentricity bias characteristic of the mature visual system.

**Figure 2.**
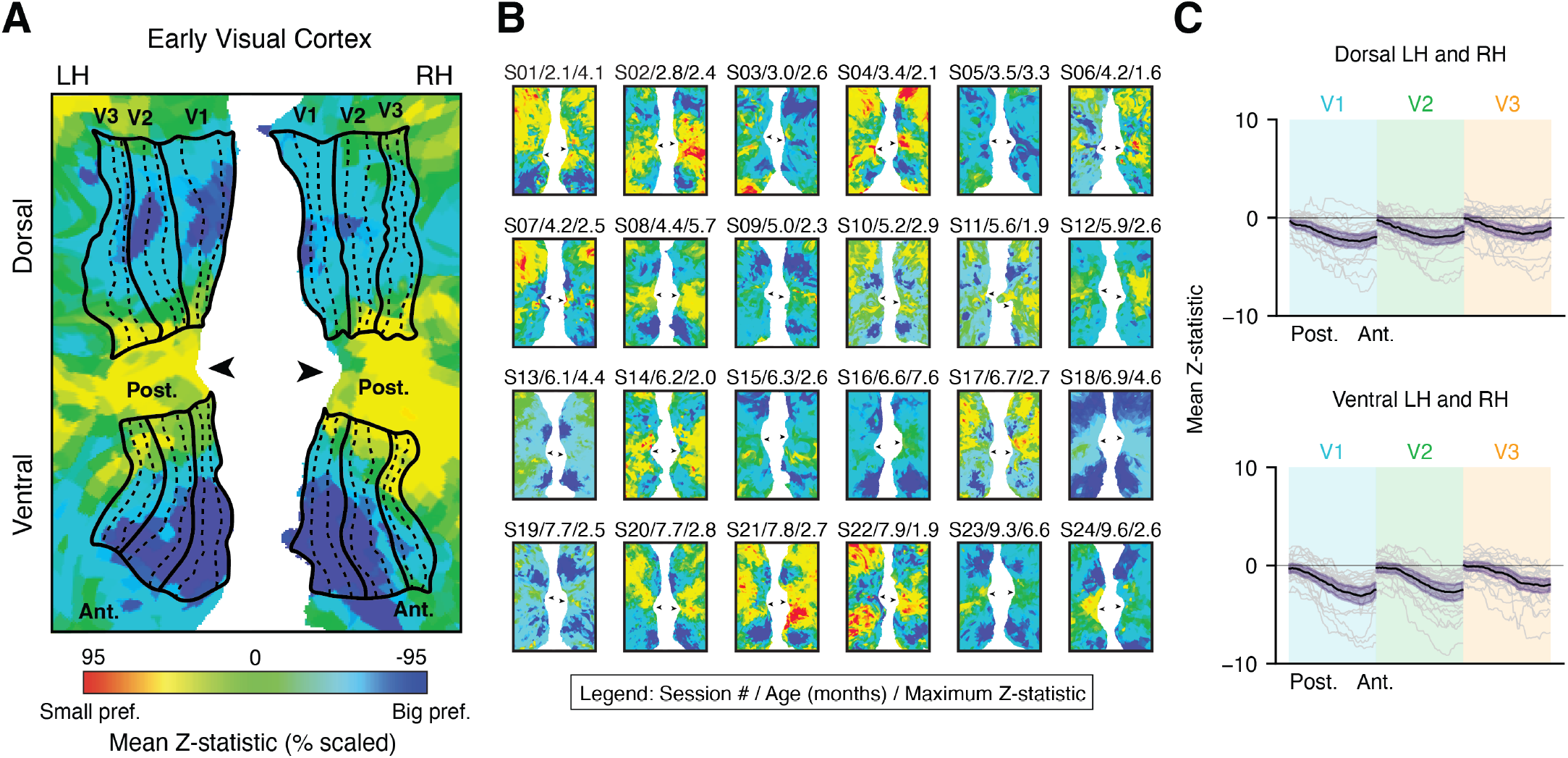
Eccentricity bias in infant early visual cortex. (A) Flattened cortical surface of an example 4.4-month-old infant overlaid with the big-versus-small contrast map. Solid lines delineate dorsal and ventral V1, V2, and V3, and dashed lines show the trajectories sampled from the posterior (post.) to anterior (ant.) axis within each visual area. Colors indicate the Z-statistic value for the big-versus-small contrast. The color scale was normalized to the 95th percentile of voxel-wise absolute Z-statistics within a standard occipital lobe parcel. Arrows indicate the location of the occipital pole along the calcarine sulcus. (B) Big-versus-small contrast maps for all infants. Labels above each surface indicate the participant’s session number, participant age in months, and maximum Z-statistic used for visualization. (C) Contrast values sampled parallel to the area boundaries were resampled from the posterior to anterior (i.e., fovea to periphery). Dorsal and ventral gradients are shown separately, collapsed across left (LH) and right hemispheres (RH). Gray lines show individual participants, the black line shows the mean across participants, and the purple-shaded region indicates the bootstrapped 95% confidence interval of the mean.

We next asked whether an eccentricity bias extends beyond EVC into ventral temporal cortex (VTC)^23^. In an exploratory analysis, we measured the size contrast along the lateral-to-medial axis (corresponding to the foveal-to-peripheral axis) in infants’ anatomically defined VTC. We found a significant increase in preference for big stimuli in medial VTC compared to lateral VTC (CI = [0.92, 1.71], p *<* 0.001), revealing an eccentricity gradient across infant VTC (Figure S3).

### Evidence for size, but not category, responses in infant ventral temporal cortex

Having established eccentricity gradients in EVC and VTC, we next asked whether the collateral sulcus, defined using a probabilistic functional atlas^24^, responds more strongly to size (big-versus-small contrast) or category (scenes-versus-letters contrast). We hypothesized that the size effect would be present in CoS throughout infancy, while a category effect would only emerge in older infants (6–10 months).

In our adult sample, CoS showed both effects: a significant response to size (mean Z-statistic = 3.165, Cohen’s d = 1.821, p = 0.017) and category (mean Z = 4.077, d = 2.337, p = 0.017), confirming that our paradigm elicited the expected pattern of both eccentricity and category effects in the CoS of adults (Figure S4). In exploratory analyses, we compared the magnitude of the size contrast with the magnitude of the category contrast and found no difference between them (mean Z = -0.682, d = -0.486, p = 0.162).

In infants, we found a significant effect for big versus small stimuli (mean Z = 0.932, d = 0.638, p = 0.003), consistent with our preregistered prediction that CoS would show a greater response to big than small stimuli, regardless of age or category (Figure 3). We found no significant preference for scenes over letters, either across the full sample (mean Z = 0.112, d = 0.094, p = 0.654), or in the 6-10 month olds (N = 12; mean Z = 0.080, d = 0.061, p = 0.839). Indeed, we found a non-significant relationship between age and category (r = -0.062, p = 0.778), inconsistent with our prediction that a scene preference would emerge in older infants. However, consistent with our hypothesis, we found that the size contrast was significantly larger than the category contrast (mean Z = 0.622, d = 0.471, p = 0.031). Thus, we found evidence that eccentricity is a more established property of infant ventral temporal cortex than category-selectivity.

**Figure 3.**
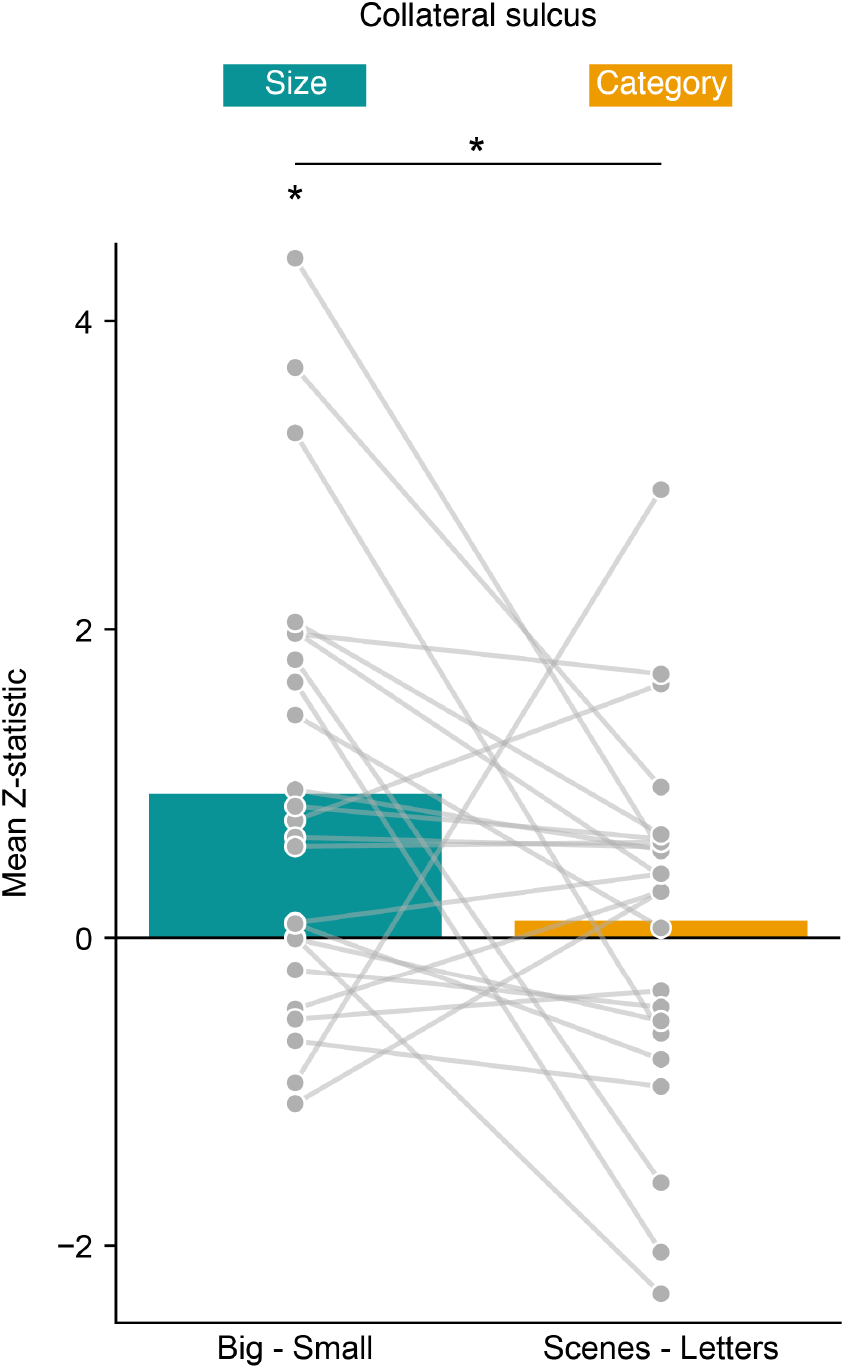
Size and category contrasts in infant ventral temporal cortex. Mean big-versus-small and scenes-versus-letters contrast values in the collateral sulcus (CoS), averaged across the left and right hemispheres. Infant CoS showed a significant size effect but no significant category effect, and the magnitude of the size contrast was significantly greater than that of the category contrast. Gray points and connecting lines show individual participants; bars show the group mean. Contrast values are mean Z-statistics, reflecting the strength of each effect relative to its estimated noise. The horizontal line indicates the significant difference between the magnitudes of the size and category contrasts. Asterisks indicate p *<* 0.05.

It is notable that we did not observe a scene preference, yet other infant studies have^17,18^. Possible reasons for this discrepancy include differences in the amount of data collected, ROI definition, and stimulus design. We found that restricting our sample to a subset of participants (N = 6) with more data showed a significant size effect (mean Z = 1.318, d = 0.788, p = 0.031) but no category effect (mean Z = 0.216, d = 0.163, p = 0.685; Figure S8). To examine the contribution of ROI definition, we conducted exploratory analyses in a subset of infants who completed an independent functional localizer. We defined subject-specific scene-selective ROIs using a scenes-versus-objects contrast. Within these ROIs, the scenes-versus-letters contrast was not significant (N = 10; mean Z = -0.192, d = -0.135, p = 0.666). The size contrast was numerically larger than the category contrast but also did not reach significance (mean Z = 0.688, d = 0.374, p = 0.276; Figure S5). Moreover, the scenes-versus-letters effect was significantly weaker in the subject-specific functional ROI than in the probabilistic ROI (mean difference = -0.250, d = -1.304, p = 0.007). The reason for this result may be that the features distinguishing scenes from objects are not the same features that distinguish scenes from letters in infancy.

In a separate exploratory analysis across the full infant sample, we tested whether the category effect depended on stimulus size. When category was contrasted using only the big stimuli (i.e., big scenes vs. big letters), infants showed a significant scene preference (mean Z = 0.416, d = 0.474, p = 0.030). However, equally-powered size contrasts were numerically larger (big scenes vs. small scenes: mean Z = 0.957, d = 0.723, p = 0.001; big scenes vs. small letters: mean Z = 0.767, d = 0.575, p = 0.011; Figure S6). Across these exploratory analyses, we find that while the category effect can be increased, the size effect remains stronger.

As a control region, occipital temporal sulcus was not expected to show a size effect in either group, since the fovea was stimulated in both the big and small conditions. Consistent with this, neither adults (mean Z = 0.045, d = 0.022, p = 0.938) nor infants (mean Z = -0.122, d = -0.103, p = 0.624) showed a significant preference for small over big stimuli. For category, adults showed a significant preference for letters over scenes (mean Z = 3.779, d = 4.318, p = 0.016). As expected, the pre-reading infants showed no such letter preference (mean Z = 0.113, p = 0.618; Figure S7).

## DISCUSSION

We found that an eccentricity bias is present throughout infant visual cortex and is prominent in regions that become category-selective in adults. Responses to big versus small stimuli showed a systematic gradient along the posterior-to-anterior axis in areas V1-V3 of early visual cortex and along the lateral-to-medial axis of ventral temporal cortex. In the collateral sulcus, which is scene-selective and peripherally-biased in adults^5–7^, infants responded more strongly to stimuli extending into the periphery, yet showed no reliable category preference for scenes over letters. These results suggest a developmental sequence in which an eccentricity bias is evident before a reliable category preference emerges.

The pattern observed here is consistent with the proto-organization hypothesis^9,21^. On this account, low-level features first organize the developing visual system, providing a scaffold for subsequent experience-dependent refinement. Eccentricity maps may be one such organizing feature. Different categories place distinct demands on visual processing because the information needed to recognize them is distributed differently across the visual field. For scenes, critical information about the environment extends beyond the center of gaze: extracting the overall structure of an environment to guide actions and eye movements requires integrating information from the periphery. Acquiring competence with scenes may therefore involve refining responses in regions that are already biased toward peripheral input, contributing to the localization of scene-selective cortex^7^. By contrast, the information that distinguishes letters falls within fixation, so foveally-biased regions may be better positioned to become letter-selective. In this way, an eccentricity bias could provide an initial scaffold for the development of category-selective cortex.

Our findings extend work on infant retinotopy in EVC^15^ by showing that eccentricity can be measured in infants without requiring strict fixation. This opens new possibilities for studying the development of retinotopic organization at ages where conventional mapping paradigms are not feasible. Here, a task-based manipulation of stimulus size revealed systematic eccentricity gradients across V1-V3 in infants as young as 2 months, the earliest evidence of task-based eccentricity organization in primates, consistent with earlier findings using functional connectivity^9,23^. The eccentricity gradients observed here align with the spatial frequency gradients we previously observed in infant visual cortex^15^, demonstrating that multiple overlapping dimensions of visual organization are present in infancy^9–14^.

Although an eccentricity bias was evident, the preference for scenes over letters in infant CoS was significantly weaker than the response to stimulus size. This suggests that the category-selectivity observed in previous studies^17,18^ is not the primary organizing property of infant ventral temporal cortex. Notably, while the category effect was not significant in our preregistered analyses, an exploratory comparison revealed a significant preference for scenes when both categories were presented at the larger size (40°). One possible explanation is that infants’ limited visual acuity made scenes and letters harder to distinguish at the small size (6°). Contrary to this explanation, category preference did not increase with age, as might be expected with improvements in acuity. Instead, larger stimuli may elicit stronger responses in scene-selective cortex, increasing sensitivity to differences between categories, as has been shown in adults^25^. Moreover, defining scene-selective voxels using each participant’s own data from an independent task did not strengthen the category contrast measured in the present task. Future work is needed to investigate whether the features that distinguish scenes from objects differ from those that distinguish scenes from letters in the infant brain. If so, the effort needed to observe scene-selectivity is, in itself, an indicator that eccentricity is more established in infancy.

In conclusion, we used fMRI with awake, behaving infants to show that an eccentricity bias is present across infant visual cortex, including both early visual areas and higher-level ventral temporal cortex. Our work demonstrates the feasibility of studying retinotopic organization throughout infant visual cortex. More broadly, our results suggest that where information falls in the visual field may be foundational to the developing visual system, shaping the emergence of later functional specialization.

## METHODS

### Participants

We preregistered a sample of 24 sessions, with 12 infants aged (2-6 months) and 12 aged (6-10 months) to ensure balanced coverage across 2-10 months. Our final infant sample included 22 unique participants, aged 2.1-9.6 months (M = 5.75 months, SD = 2.03; 14 females) (Table 1). Caregiver-reported race was American Indian or Alaska Native (N = 1), Asian (N = 9), multiracial (N = 2), and White (N = 10). Two participants contributed two sessions each, one in each age group, and these were treated as independent observations in line with previous research^16^,^15^. An additional 18 sessions were attempted but were excluded from the final sample because they did not meet the inclusion criteria for sufficient usable functional data (N = 10), did not have a usable anatomical image for surface reconstruction (N = 7), or had two usable sessions within the same age group, only one of which we could include (N = 1).

**Table 1.**
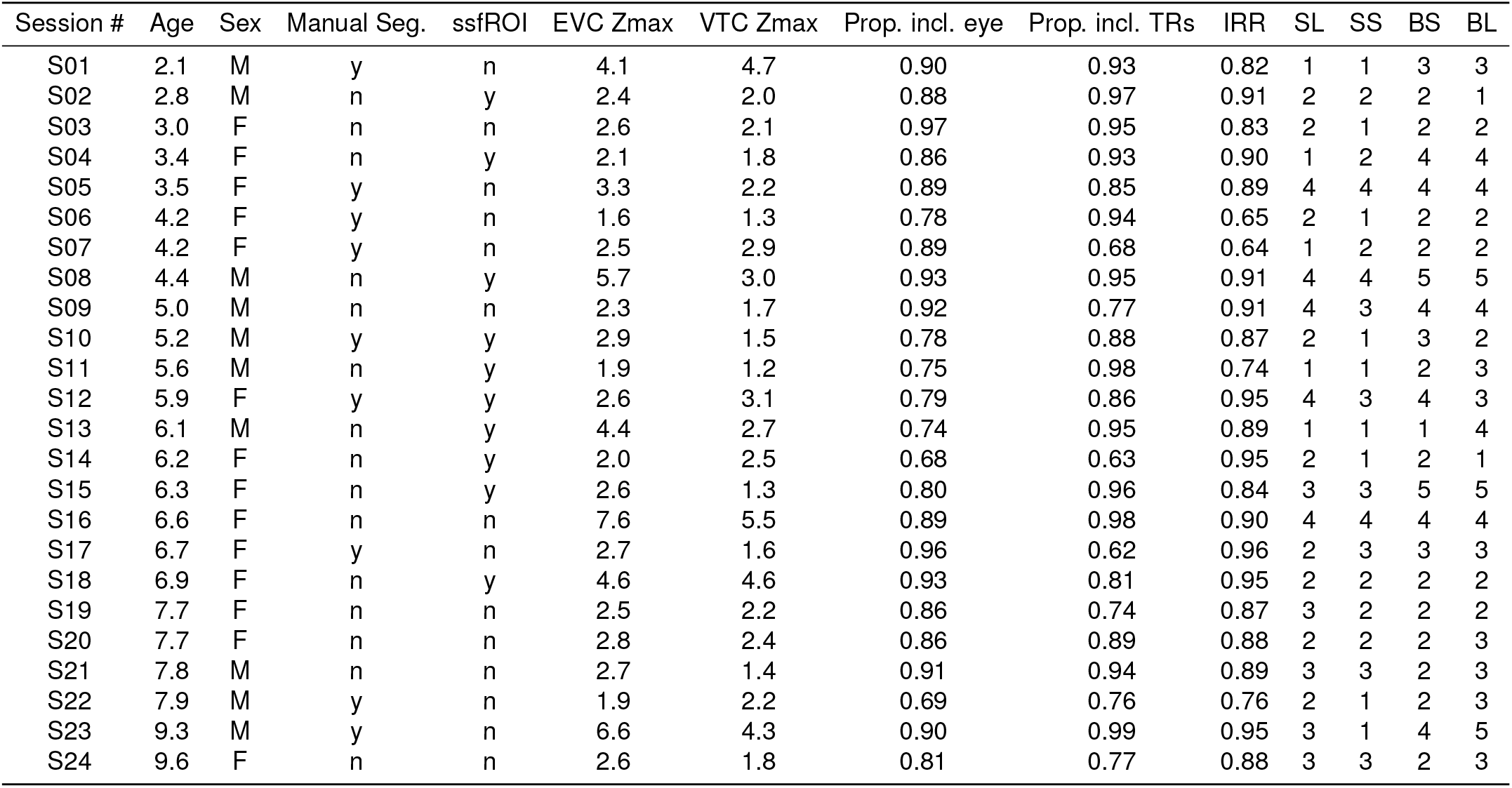
Participant information. Information for included infant participants. ‘Session #’ is each participant’s session ID. ‘Age’ is reported in months. ‘Sex’ is female or male. ‘Manual Seg.’ is whether the participant’s segmentation was manually edited. ‘ssfROI’ is whether they had a subject-specific functional ROI. ‘EVC Zmax’ is the 95th percentile of voxel-wise absolute Z-statistics within a standard occipital lobe parcel. ‘VTC Zmax’ is the 95th percentile of voxel-wise absolute Z-statistics within a combined CoS and OTS standard parcel. ‘Prop. incl. TRs’ is the proportion of usable TRs from the included blocks. ‘Prop. incl. eye’ is the proportion of usable eye data. ‘IRR’ is the interrater reliability for coding total on-screen gaze (collapsing across center, off-center, and undetected responses). ‘SL’,’SS’,’BS’, and ‘BL’ are the number of blocks per condition for small letters, small scenes, big scenes, and big letters, respectively.

An adult sample (N = 7, 3 females; M = 23.59 years, SD = 2.90) was collected before preregistration to validate our paradigm.

Participants were recruited through fliers, through promoted social media advertisements or word of mouth. This study was approved by the Institutional Review Board at Stanford University. Adults provided written informed consent for themselves (if they were the participants) or on behalf of their child (if their child was the participant).

### Data acquisition

Data were acquired with a 3T GE UHP MRI scanner at the Center for Cognitive and Neurobiological Imaging at Stanford University, using a custom 32-channel infant head coil^26^. Functional images were collected with a whole brain T2* echo-planar imaging sequence (infant participants: TR=2000ms, echo time=33ms, flip angle=71, matrix=68x68, slices=33, resolution=3mm isotropic; adult participants: same as above except slices=35). Anatomical images were acquired using one or more of the following sequences, depending on the infant’s wakefulness and age: T1-weighted: TR = 2843 ms, TE = 2.8 ms, flip angle = 8°, matrix = 256 × 256, 140 slices, voxel size = 1 mm isotropic. T1-weighted: TR = 7.8 ms, TE = 3.236 ms, flip angle = 12°, matrix = 256 × 256, 158–166 slices, voxel size = 1 mm isotropic. T2-weighted: TR = 3648 ms, TE = 101.7 ms, flip angle = 90°, matrix = 256 × 256, 132 slices, voxel size = 1 mm isotropic. B0 fieldmaps were acquired using a dual-echo gradient echo sequence (TR = 100 ms, TE = 3.8, 6.3 ms, flip angle = 20°, matrix = 64 × 64, 44 slices, voxel size = 3.4 mm isotropic).

### Procedure

A variant of our approach for collecting fMRI data from awake infants has been described in a previous methods paper^27^, with important details and customizations described below. Families visited the lab before their first scanning session for an orientation session, aimed at acclimating the infant and parent to the scanning environment. Scanning sessions were scheduled for a time when parents expected their infant to be calm and awake. Participants were extensively screened for any metal on or in their body. Infants were then fitted with hearing protection, including silicone inner-ear putty and noise-canceling headphones (OptoACTIVE Tiny™; OptoAcoustics, Ltd). After, the infant was positioned in a custom head coil^26^ with their body on top of a bean bag. Once in the scanner, stimuli were presented on a custom-ceiling mounted screen via a mirror-based projection system. A video camera (MRC 12m-i) recorded the infant’s face via a peephole in the ceiling screen to monitor their state and for offline gaze coding. Adult participants underwent the same procedure with the following exceptions: they did not attend an orientation session, they had ear plugs and noise-canceling headphones (OptoACTIVE Slender™; OptoAcoustics, Ltd), they were not on a bean bag, and they used the Nova Medical 32-Channel Head Coil.

Experimental stimuli were shown using Psychtoolbox^28^ for MATLAB. For the infant participants, multiple experiments were sometimes run within the same scanning session. If more than one experiment occurred within the same functional run, the run was divided into experiment-specific pseudo-runs for subsequent preprocessing and analyses. For this paper, we will focus on one primary experiment run in all participants reported here, as well as a second experiment (i.e., a category localizer) that was run in a subset of participants reported here.

### Study Design

The experiment used a 2 × 2 factorial design that varied stimulus category (scenes vs. letters) and size (big vs. small), creating four stimulus conditions: big scenes, small scenes, big letters, and small letters. Each condition was presented as a block, lasting 14 s. Within each block, stimuli of the same condition were presented at 1 Hz (700 ms on, 300 ms off). After every set of four conditions, there was a blank period lasting 6 s. All stimuli were presented in a circular aperture and differed in size and content. Big stimuli were 40°in diameter, while small stimuli were 6°. Scene stimuli consisted of outdoor natural environments, excluding city scapes and indoor spaces^29^. Letter stimuli were generated by overlaying different segments of a Latin paragraph onto phase-scrambled versions of the scene images. A contrast of scenes versus phase-scrambled scenes has been sufficient to identify scene-selective cortex in adults^25^. The images within each condition were yoked across stimulus category. For instance, the sequence of scene images that were shown in a block were repeated as phase-scrambled backgrounds for the letter stimuli, with different sequences occurring for big and small stimuli.

### Gaze Coding

Infant gaze was manually coded offline by two or three coders (M = 2.5) who were blind to the condition being coded. Coders classified each frame as either center (e.g., eyes oriented on the center of the stimulus), off-center (e.g., eyes oriented to the side of the stimulus), off-screen (i.e., blinking or looking away), or undetected (e.g., the eyes are occluded by the infant’s arm).

We computed inter-rater reliability (IRR) as each coder’s mean frame-by-frame agreement with the other coder. If IRR was below 0.7 (across all four label types), a third coder rated the session. The coder with the lowest mean reliability was excluded, and the remaining two coders were used in subsequent analyses. IRR for coding across all four label types was moderate (M = 0.744; SD = 0.095; range = 0.54-0.93). IRR for total on-screen gaze (collapsing across center, off-center, and undetected responses) was high (M = 0.86; SD = 0.088; range = 0.64-0.96).

To determine block inclusion, we assigned each frame a gaze label based on the modal response across both coders within a five-frame window centered on that frame. Blocks were excluded if the resulting gaze estimate indicated that the infant was off-screen for more than 50% of the block.

To test for differences in looking behavior across size and category, we computed each participant’s mean proportion of on-screen frames coded as center in each condition. We then used permutation testing to compare center-looking between big and small stimuli and between scene and letter stimuli.

### Preprocessing

Data were analyzed using a custom pipeline designed for awake infant fMRI preprocessing^27^ and surface reconstruction^15^. At least three burn-in volumes were discarded from the start of each run. We performed motion correction on each run, aligning to the centroid volume (i.e., the volume that minimizes the Euclidean distance to all other volumes). Time points with greater than 3mm of translational motion were temporally interpolated in order to minimize their effect on preprocessing. These time points were also excluded in subsequent analyses. Slices were realigned using slice-time correction. The brain was masked, smoothed using a 5mm Gaussian kernel, and linearly detrended. Despiking was used to remove aberrant time points within voxels^30^. After removing excess time points, the data were Z-scored in time within a run. Using the FMRIB Software Library (FSL), fieldmap correction was performed to correct for the B0 distortion, and aligned the centroid to the anatomical (6 degrees of freedom). The alignment between the anatomical and standard space (MNI152) was achieved using Advanced Normalization Tools (ANTs) in two steps. First, each participant’s anatomical image was nonlinearly registered to an age-appropriate infant template based on the participant’s age at scan. Second, a predefined non-linear transformation was used to align the infant template to the adult MNI152 standard. These two transformations were performed to put all functional data in standard space, allowing comparisons to be made across participants.

For surface reconstruction, we used the BIBSNet Docker container^31^ to generate gray and white matter tissue segmentations from the T1- or T2-weighted images. We manually corrected these BIBSNet segmentations (N = 9) to fix errors in the white and gray matter labels (e.g., mislabeled white matter voxels) using ITK-SNAP. The corrected segmentation replaced the segmentations used by infant FreeSurfer^32^, producing a mesh of each infant’s cortical surface. To generate cortical flat maps, we cut the inflated surface into patches, then flattened using the standard FreeSurfer procedure. Functional data were resampled in SUMA^33^ using an icosahedral mesh, aligning each participant’s surface to a standardized spherical space with a constant number of nodes.

We used the functional atlas of visual cortex^24^ in MNI standard space to define the CoS and OTS ROIs. The CoS ROI was created by combining the left and right hemisphere CoS ROIs (0% probability threshold) from their atlas. The OTS ROI was defined by the left inferior occipital sulcus (IOS) and posterior occipitotemporal sulcus (pOTS) ROIs (0% probability threshold). The EVC ROI was defined using the ventral and dorsal V1 (0% probability threshold) from an adult-defined atlas^22^.

### GLM analysis

The functional data were analyzed using a General Linear Model (GLM) implemented in FEAT (fMRI Expert Analysis Tool), part of the FSL (FMRIB Software Library), where a GLM was fit to the BOLD activity in each voxel. Each regressor modeled one of the four stimulus conditions with a boxcar lasting 14 s, convolved with a double-gamma hemodynamic response function. The six translation and rotation parameters from motion correction were included in the GLM as nuisance regressors. TRs that were excluded (i.e., had translational motion greater than 3mm) were scrubbed by including an additional regressor for each to-be-excluded time point^34^. Contrasts were defined in FEAT to compare the four experimental conditions (small letters, small scenes, big letters, and big scenes). We extracted Z-statistics for each participant and contrast, then computed the mean Z-statistic within each ROI.

In an exploratory analysis, we computed the difference between the main effect of size (big vs. small) and the main effect of category (scenes vs. letters). The two contrasts can be written out as follows:

(big scenes + big letters) - (small scenes + small letters) [size contrast].

(big scenes + small scenes) - (big letters + small letters) [category contrast].

Two of the conditions (big scenes and small letters) have the same sign across these two contrasts, so subtracting these contrasts cancels them out. The remaining two conditions (big letters and small scenes) have different signs in the two contrasts, so to compute the difference of the contrasts, we simply contrasted big letters and small scenes.

To evaluate group-level effects, we used permutation testing with 10,000 randomizations to determine whether the mean Z-statistics across sessions were significantly different from zero within each ROI. A response was considered significant if the proportion of permutations resulting in a mean Z-statistic greater than or equal to the observed mean was sufficiently small, as determined by the p-value.

### Gradient Tracing

We overlaid early visual areas V1 through V3 from the Wang atlas (Wang et al., 2015) onto the cortical surface of each participant’s left and right hemisphere. Following a previous approach^15^, we drew two equally spaced lines running posterior to anterior in each area, corresponding to the foveal-to-peripheral axis of each visual area. Lines were drawn in six areas total: ventral V1, V2, and V3, and dorsal V1, V2, and V3. Lines were drawn using only the V1 through V3 boundaries for guidance, with no contrast maps visible, so placement was blind to the underlying data (i.e., to local intensity changes within each area).

For each line, Z-statistic values from the big-versus-small contrast were extracted along the line and resampled to a fixed number of points (60 per area) to account for differences in line length across participants and hemispheres. The two lines within each area were averaged together, and eccentricity profiles were then averaged across the left and right hemispheres. To visualize the contrast maps, the color scale for each participant was set using the 95th percentile of voxel wise Z-statistics within a standard occipital lobe parcel.

To test whether the big-versus-small effect changed systematically within an area, we compared the mean response in the first quartile (foveal end) of each area’s profile to the mean response in the last quartile (peripheral end) using permutation testing for significance and bootstrap resampling for confidence intervals. To compute the bootstrapped CIs, we performed 10,000 resamples in which sessions were sampled with replacement, and the mean difference was recomputed. 95% confidence intervals were taken as the 2.5th and 97.5th percentiles of the resampled distribution.

To trace the eccentricity gradients in ventral temporal cortex (VTC), we followed previous approaches^35–37^. Specifically, we defined each infant’s VTC anatomically, using the occipital temporal sulcus (OTS), posterior transverse collateral sulcus (ptCoS), collateral sulcus (CoS), and the anterior tip of the mid-fusiform sulcus (MFS), which aligns with the posterior end of the hippocampus. These region boundaries were identified using Freesurfer’s anatomical annotation file (aparc.annot). After defining the VTC boundaries, we drew 7 lines equally spaced going from the lateral-to-medial axis. As above, the contrast maps were not visible when these lines were drawn. Values along these lines were averaged to test for an eccentricity gradient across VTC. To visualize the contrast maps, the color scale for each participant was set using the 95th percentile of voxel wise absolute Z-statistics within a standard parcel comprising the OTS and CoS regions from the Rosenke atlas used in the main analyses.

### Functional ROI analysis

To assess whether ROI definition influenced the effect of scenes versus letters, we conducted an exploratory analysis in a subset of participants (N = 10) who completed an independent event-related experiment containing scenes, objects, and faces (12 exemplars per categoryeach presented for 2.5 s with a 1.5–2.5s ISI and repeated twice). We used the scenes-versus-objects contrast to localize the scene response. The remaining participants either did not complete this experiment or did not meet our inclusion criterion of at least 12 usable events per category.

We defined a search space by dilating the Rosenke CoS parcel^24^ used in the main analyses to approximately match the volume of the Julian PPA parcel^38^, which is about 2.7 times larger by voxel count. Dilation was performed in MNI standard space using three iterations of binary dilation. For each participant, we defined a subject-specific functional ROI (ssfROI)^17,18^ within this search space using data from the independent experiment. The voxels were ranked by Z-statistic from the scenes-versus-objects contrast. The top 10% of voxels within the search space were retained as that participant’s ssfROI. We then extracted mean Z-statistics from the scenes-versus-letters contrast in each participant’s main-task data within their ssfROI, and tested for a significant scene preference using a permutation test.

### Deviations from preregistration

We followed the preregistration for sample size, design, and statistical tests, but made unintentional deviations in participant inclusion and an analysis specification. Sessions were included in our sample based on the following criteria: the participant *viewed* at least two repetitions of each condition, at least one repetition of each condition was *usable*, and we acquired an anatomical scan sufficient for cortical surface reconstruction. These criteria deviated from our preregistration, which required at least three *usable* repetitions per condition and did not include an anatomical-quality criterion. The repetition criteria deviation occurred at the start of data collection, in which we aimed for four blocks, only exceeding four blocks in the case of an engaged participant that had some unusable data. This goal was misaligned with the usability threshold since we assume that 50% of the data will be unusable on average. The anatomical-quality criteria deviation was necessitated by our surface-based analyses, which require a usable cortical surface reconstruction (lower quality data is adequate for volumetric alignment), but was not anticipated at the time of preregistration. Both discrepancies with our preregistration were identified only during manuscript preparation. To evaluate the effects of these deviations, we repeated our primary analyses but restricted to participants who met the original three block criterion, along with the anatomical quality threshold. Despite a smaller sample size (N = 6), the primary contrast between big and small was significant, whereas the category contrast was not (Figure S8). We also deviated from the preregistered analysis by examining early visual areas V1 through V3, rather than V1 through V4, as we chose to only analyze areas that could be sampled symmetrically across dorsal and ventral cortex.

## RESOURCE AVAILABILITY

### Lead contact

Requests for further information and resources should be directed to and will be fulfilled by the lead contact, Sarah Tung.

### Materials availability

The design for the custom projection system used for data collection is publicly available at https://soc.stanford.edu/resource/.

### Data and code availability

- The code for running the eccentricity task can be found at: https://github.com/scaffolding-of-cognition-team/soc_menu/tree/Whole_Eccentricity
- The preregistered plan for the study can be found at: https://osf.io/v8qfd/overview
- The code for performing the analyses can be found at: https://github.com/scaffolding-of-cognition-team/soc_neuropipe/tree/Whole_Eccentricity
- The raw and preprocessed data will be released on Dryad at the time of publication.

## ACKNOWLEDGMENTS

This work was funded by internal Stanford funds for C.T.E. and the National Science Foundation Graduate Research Fellowship Program DGE-2146755 to S.S.T. Thank you to A.G. Ornelas for help with making the projection system. Thank you to K. Grill-Spector, V.S. Natu, and M.J. Arcaro for thoughtful discussions about the analysis. Thank you to J.D. Yeatman for helpful discussions about the design of the experiment. Thank you to K. Cepeda-Ramos, J. Noyola, and N.A. Sahrling for help with gaze coding.

## AUTHOR CONTRIBUTIONS

C.T.E. and S.S.T. created the experiment design. S.T., E.M.C., C.T., K.S., and C.T.E. collected the data. K.S. and C.T.E. created the custom projection system. S.T. preprocessed the data. S.T. and C.T.E. analyzed the data. All authors contributed to the drafting of the manuscript.

## DECLARATION OF GENERATIVE AI AND AI-ASSISTED TECHNOLOGIES

During the preparation of this work, the first author wrote the first draft and initial code. ChatGPT and Claude were used sparingly to improve manuscript clarity, provide coding assistance during analysis, and verify the accuracy of the final code. All AI-generated suggestions were critically reviewed and edited by the author, who takes full responsibility for the content of the publication.

## SUPPLEMENTAL INFORMATION INDEX

**Figure S1.**
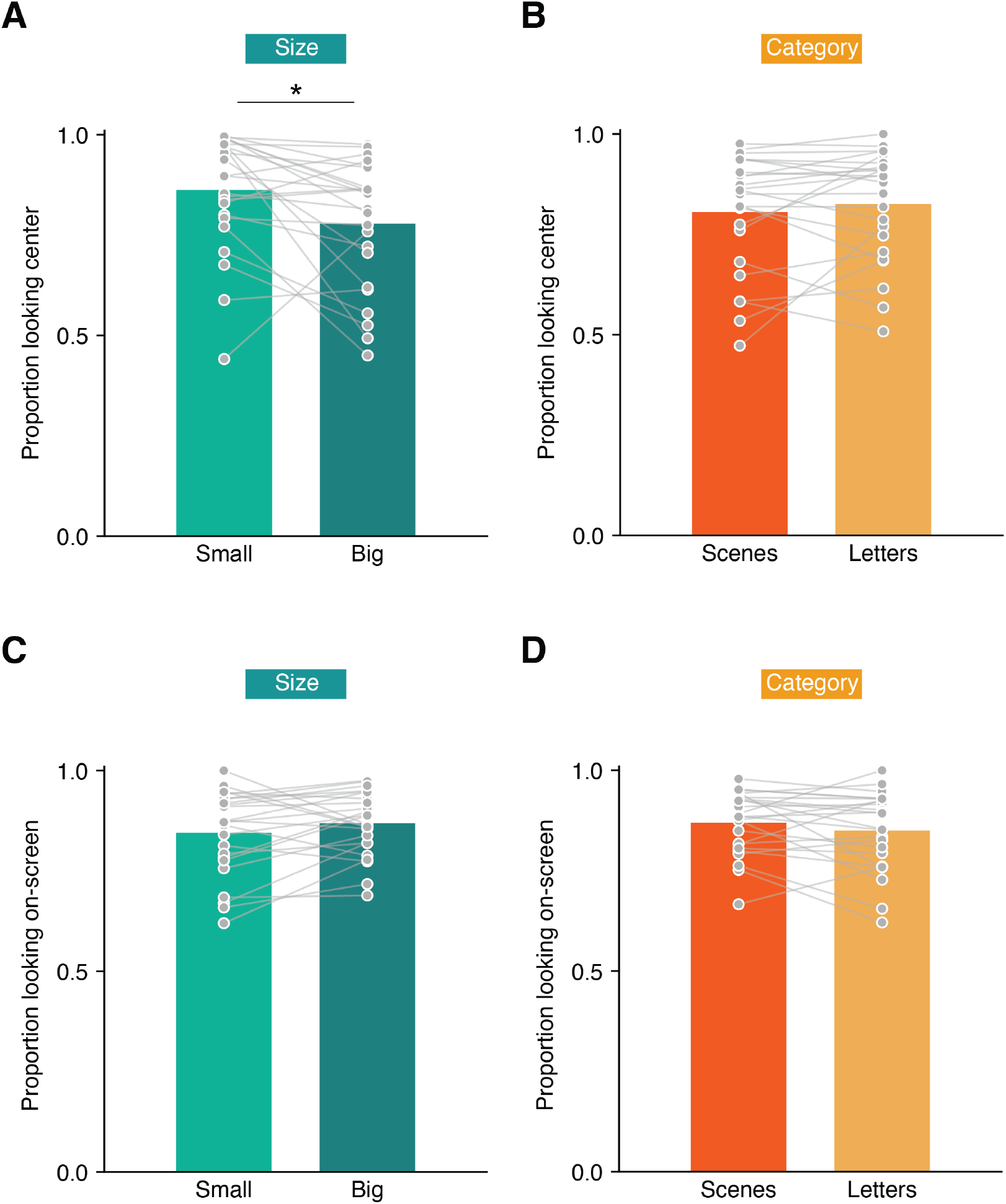
Looking behavior for size and category stimuli. (A) Proportion of looking at the center of the stimulus during small and big conditions. For each participant, center-looking was calculated as the number of center-coded frames divided by the total number of center- and off-center coded frames. Participants looked significantly longer at the center for small (M = 0.86, SD = 0.14) compared to big (M = 0.78, SD = 0.16) stimuli (mean difference = 0.08, d = 0.52, p = 0.016). Connecting lines show individual participants and bars show the group mean. Horizontal line indicates the comparison between small and big stimuli for center-looking. Asterisk indicates p *<* 0.05. (B) Same as in (A) but for the scene (M = 0.81, SD = 0.15) and letter (M = 0.83, SD = 0.14) categories (mean difference = -0.02, d = -0.21, p = 0.334) where no such difference was observed. (C) Proportion of looking on-screen during small and big conditions. For each participant, on-screen looking was calculated as the number of center-, off-center-, and undetected-coded frames divided by the total number of frames. Participants did not look longer at big (M = 0.87, SD = 0.08) compared to small (M = 0.84, SD = 0.11) stimuli (mean difference = -0.02, d = -0.32, p = 0.129). (D) Same as in (C), for scene (M = 0.87, SD = 0.08) and letter (M = 0.85, SD = 0.10) categories (mean difference = 0.02, d = 0.26, p = 0.219).

**Figure S2.**
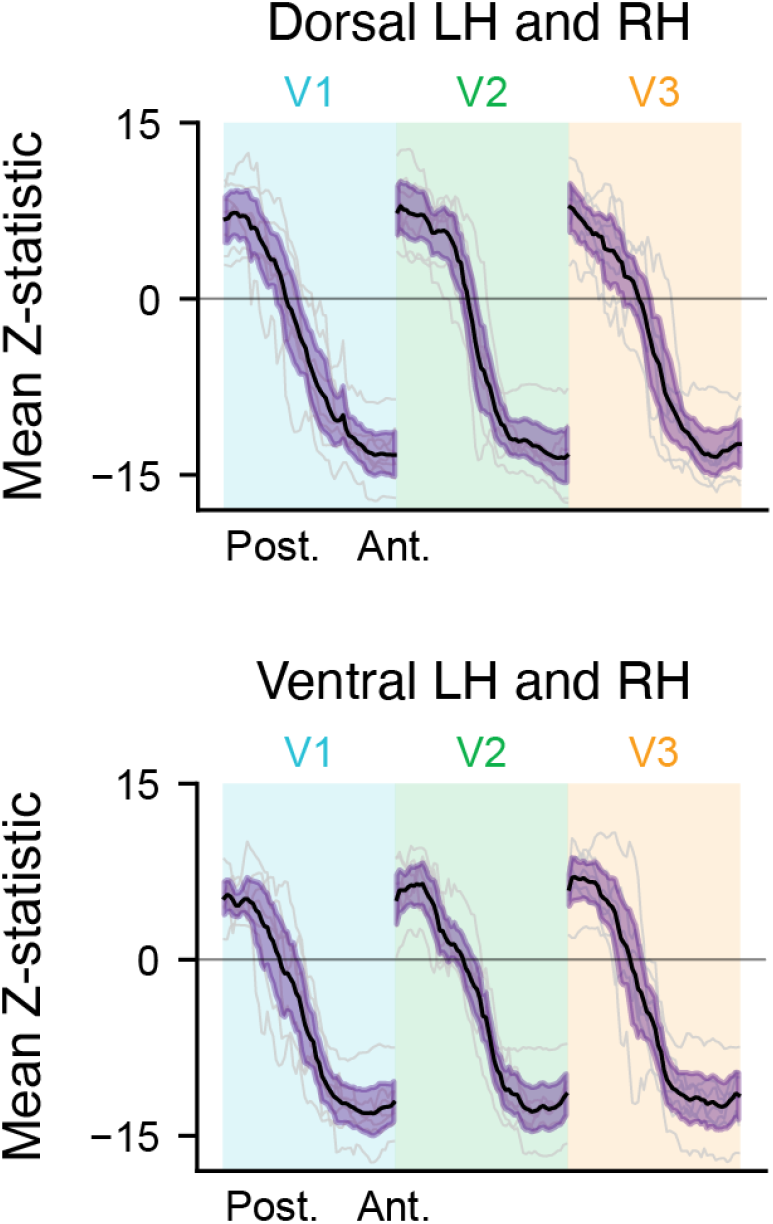
Eccentricity bias in adult early visual cortex. Big-versus-small contrast values sampled along the posterior-to-anterior axis of dorsal and ventral V1, V2, and V3, averaged across (LH) and right hemispheres (RH). Big stimuli evoked a stronger response than small stimuli in the anterior portion of all visual areas tested, relative to the posterior portion: dorsal V1 (CI = [17.30, 21.40], d = 5.83, p = 0.015), V2 (CI = [17.55, 22.09], d = 5.89, p = 0.015), and V3 (CI = [16.47, 21.23], d = 5.30, p = 0.015) and ventral V1 (CI = [15.20, 19.29], d = 5.83, p = 0.015), V2 (CI = [15.39, 20.74], d = 4.69, p = 0.015), and V3 (CI = [15.69, 20.67], d = 5.03, p = 0.015. Gray lines show individual participants, the black line shows the mean across participants, and the purple-shaded region indicates the bootstrapped 95% confidence interval of the mean.

**Figure S3.**
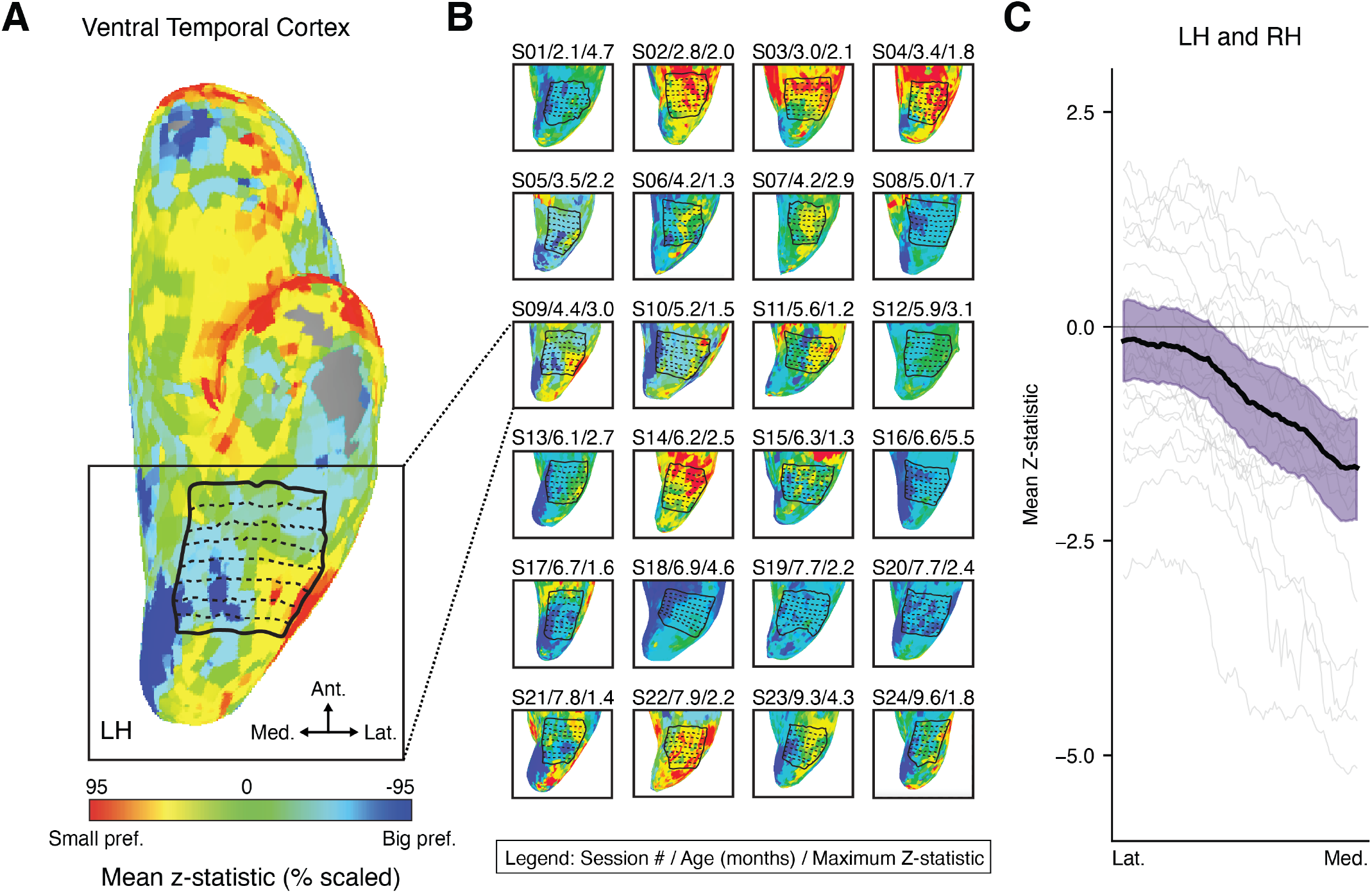
Eccentricity bias in infant ventral temporal cortex. (A) Inflated cortical surface of an example 4.4-month-old infant’s left hemisphere (LH), overlaid with the big-versus-small contrast map. Solid lines delineate the boundaries of ventral temporal cortex (VTC) and dashed lines show the trajectories sampled from the lateral (lat.) to medial (med.) axis within the VTC patch. Colors indicate the Z-statistic value for the big-versus-small contrast. The color scale was normalized to the 95th percentile of voxel-wise Z-statistics within a standard parcel comprising the occipital temporal sulcus and collateral sulcus parcel used in main analyses. (B) Big-versus-small contrast maps for all infants in the left hemisphere. Labels above each surface indicate the participant’s session number, participant age in months, and maximum Z-statistic used for visualization. (C) Contrast values sampled from the lateral to medial axis (i.e., fovea to periphery). Gray lines show individual participants averaged across their left and right hemispheres, the black line shows the mean across participants, and the purple-shaded region indicates the bootstrapped 95% confidence interval of the mean.

**Figure S4.**
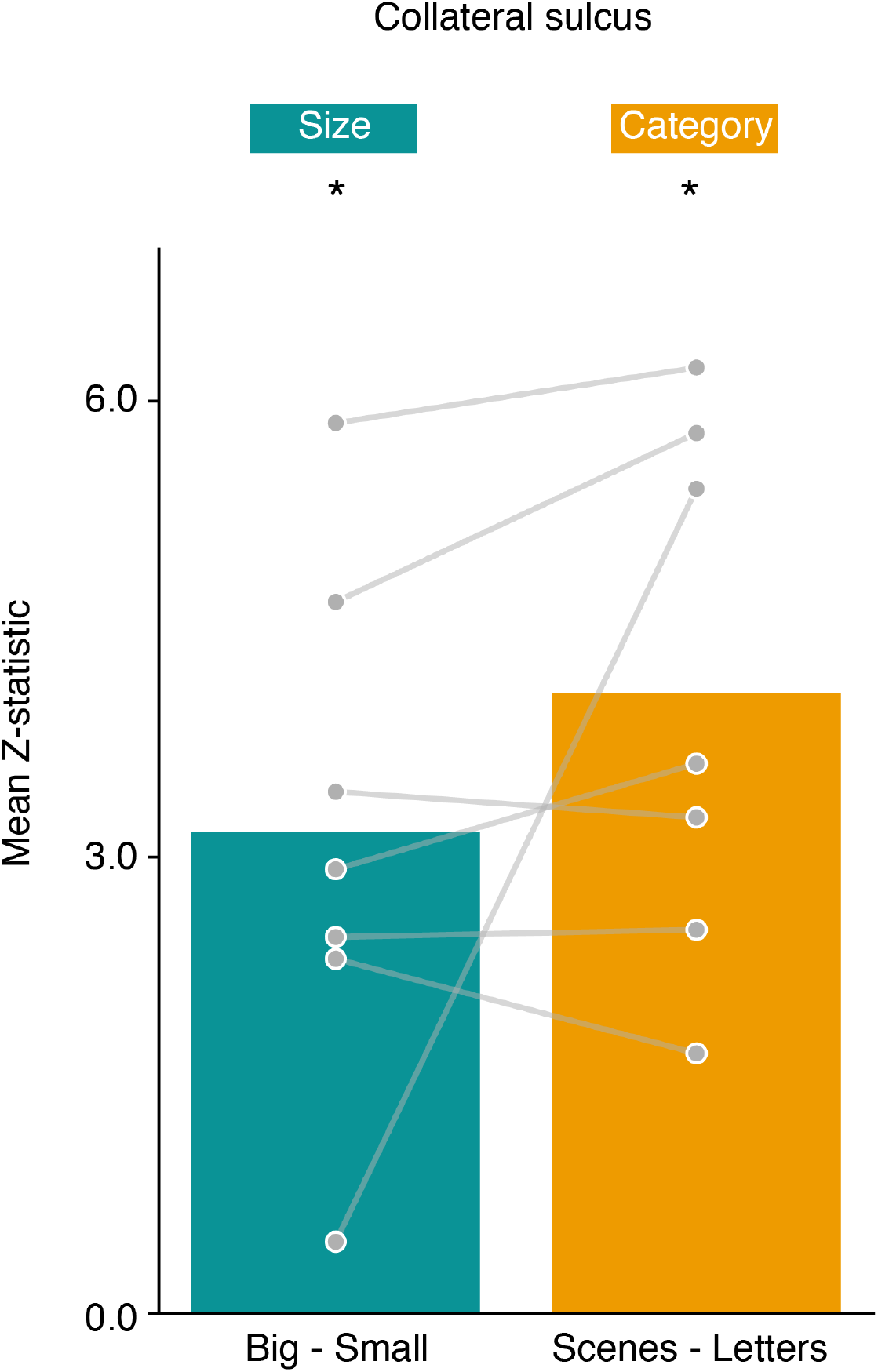
Size and category contrasts in adult ventral temporal cortex. Mean big-versus-small and scenes-versus-letters contrast values in the collateral sulcus (CoS), averaged across the left and right hemisphere. Gray points and connecting lines show individual participants; bars show the group mean. Contrast values are mean Z-statistics, reflecting the strength of each effect relative to its estimated noise. Adult CoS showed both a significant size effect and significant category effect. Asterisk indicates p *<* 0.05.

**Figure S5.**
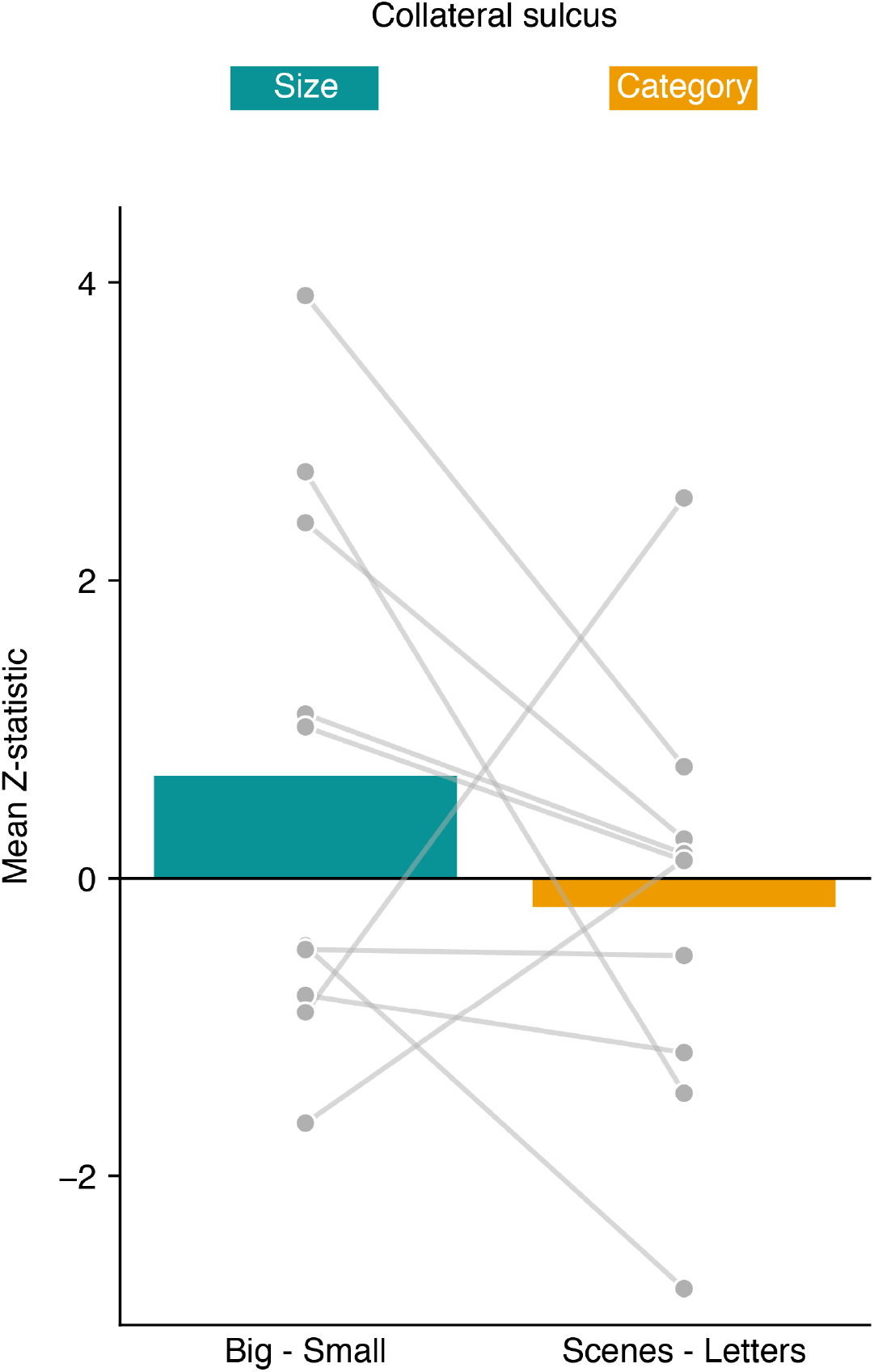
Size and category contrasts in infant ventral temporal cortex using subject-specific functional ROIs. Mean big-versus-small and scenes-versus-letters contrast values in subject-specific functional ROIs (ssfROI), defined from the scenes-versus-objects contrast in an independent localizer experiment completed by a subset of participants. Gray points and connecting lines show individual participants; bars show the group mean. Contrast values are mean Z-statistics, reflecting the strength of each effect relative to its estimated noise. Using a ssfROI did not yield a significant size or category effect.

**Figure S6.**
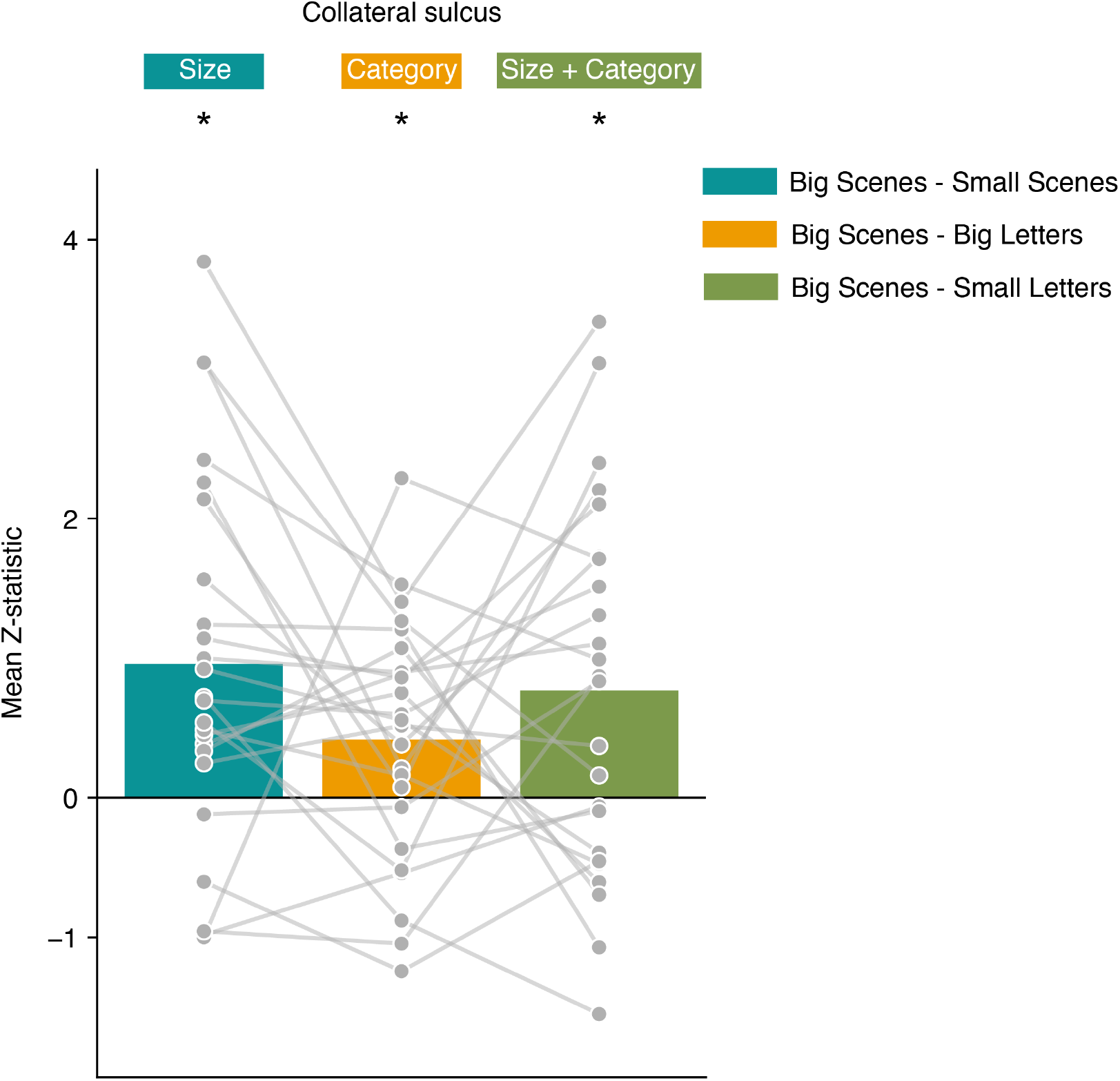
Size and category effects across matched contrasts in infant ventral temporal cortex. Mean contrast values in the collateral sulcus (CoS), averaged across the left and right hemisphere, for three comparisons: big scenes versus small scenes (mean Z = 0.957, d = 0.723, p = 0.001), big scenes versus big letters (mean Z = 0.416, d = 0.474, p = 0.030), and big scenes versus small letters (mean Z = 0.767, d = 0.575, p = 0.011). Gray points and connecting lines show individual participants; bars show the group mean. Contrast values are mean Z-statistics, reflecting the strength of each effect relative to its estimated noise. Asterisk indicates p *<* 0.05.

**Figure S7.**
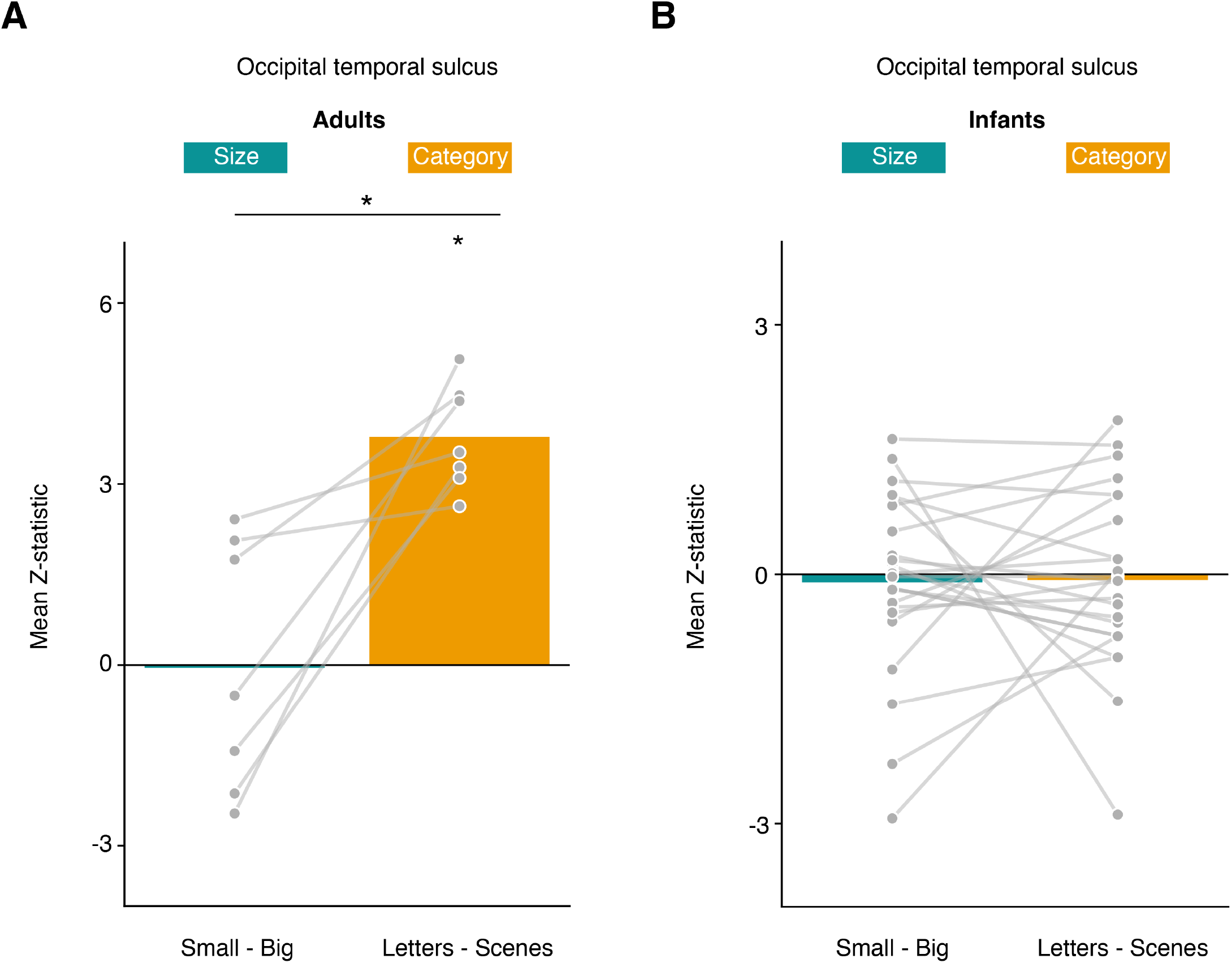
Size and category contrasts in occipital temporal cortex. (A) Mean small-versus-big and letters-versus-scenes contrast values in the occipital temporal sulcus (OTS), averaged across the left and right hemispheres in adults. Gray points and connecting lines show individual participants; bars show the group mean. Contrast values are mean Z-statistics, reflecting the strength of each effect relative to its estimated noise. (B) Same as in (A), for infants. Infant OTS did not show a significant size or category effect. Asterisk indicates p *<* 0.05.

**Figure S8.**
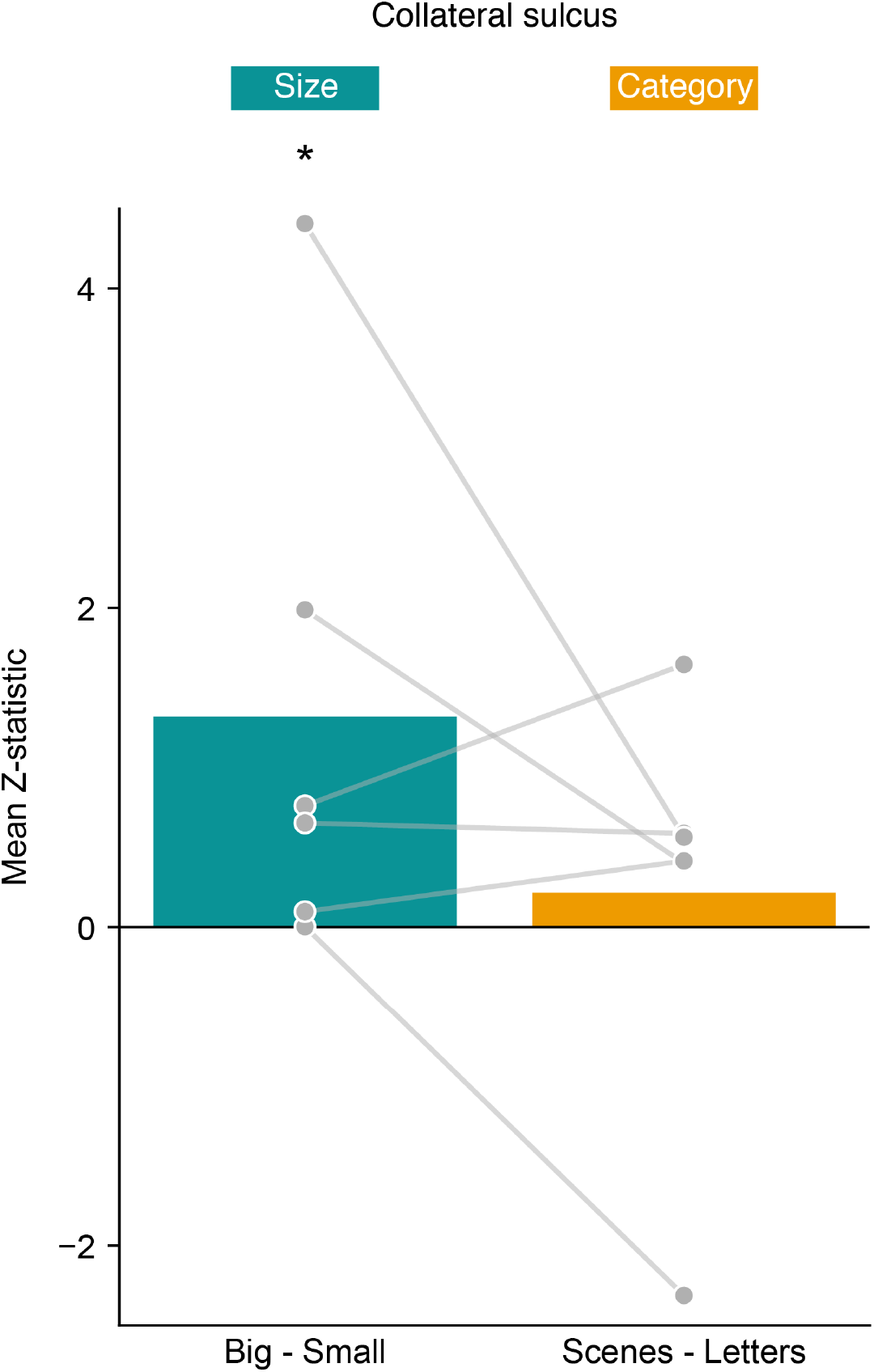
Size and category contrasts in ventral temporal cortex for infants meeting the preregistered inclusion criteria. Mean big-versus-small and scenes-versus-letters contrast values in the collateral sulcus (CoS), averaged across the left and right hemispheres. Gray points and connecting lines show individual participants; bars show the group mean. Contrast values are mean Z-statistics, reflecting the strength of each effect relative to its estimated noise. Asterisk indicates p *<* 0.05.

